# Systems Neuroscience Computing in Python (SyNCoPy): A Python Package for Large-scale Analysis of Electrophysiological Data

**DOI:** 10.1101/2024.04.15.589590

**Authors:** Gregor Mönke, Tim Schäfer, Mohsen Parto-Dezfouli, Diljit Singh Kajal, Stefan Fürtinger, Joscha Tapani Schmiedt, Pascal Fries

## Abstract

We introduce an open-source Python package for the analysis of large-scale electrophysiological data called SyNCoPy, for Systems Neuroscience Computing in Python. The package includes signal processing analyses across time (e.g. time-lock analysis), frequency (e.g. power spectrum), and connectivity (e.g. coherence) domains. It enables user-friendly data analysis on both laptop-based and high-performance computing systems. SyNCoPy is designed to facilitate trial-parallel workflows (parallel processing of trials) making it an ideal tool for large-scale analysis of electrophysiological data. Based on parallel processing of trials, the software can support very large-scale datasets via innovative out-of-core computation techniques. It also provides seamless interoperability with other standard software packages through a range of file format importers and exporters and open file formats. The naming of the user functions closely follows the well-established FieldTrip framework, which is an open-source Matlab toolbox for advanced analysis of electrophysiological data.

## Introduction

In neuroscience, methods like electroencephalography (EEG), magnetoencephalography (MEG), electrocorticography (ECoG) and microelectrode recordings are used to measure electromagnetic signals originating from brain activity. Researchers are typically interested in identifying brain activity related to certain experimental conditions, e.g., the onset of a stimulus presented to a subject. Therefore, experimental tasks are repeated many times, and the resulting trials are later averaged to reduce noise and variance. The trial repetitions combined with modern experimental setups using an increasing number of recording sites (channels), and high sampling rates can lead to very large (> 10 GB) datasets. With these datasets, standard algorithms like all-to-all connectivity computations between channels can become impossible to carry out on laptops or desktop computers with limited memory, and require workstations or high-performance computing (HPC) systems which can be complex to work with. Moreover, recently, there has been a significant surge of interest in using the scientific Python tech-stack as an open-source environment for data analysis.

Here, we present SyNCoPy (**Sy**stems **N**euroscience **Co**mputing in **Py**thon), a Python package for the analysis of large-scale electrophysiology data that combines an easy-to-use, FieldTrip-like (Oostenveld et al., 2011) application programming interface (API) with inbuilt support for distributed workflows on HPC systems.

### Related Software Packages

Scientific software packages for the analysis of neuro-electromagnetic data include FieldTrip, EEGLab (Delorme et al., 2011; Delorme & Makeig, 2004) and Brainstorm (Tadel et al., 2011) for Matlab, and MNE Python (Gramfort et al., 2013, 2014) and Elephant (Denker et al., 2023) for Python.

FieldTrip is a Matlab toolbox that was first published in 2011 and has been actively evolving since then. Its features include pre-processing, multivariate time-series and connectivity analysis and source localization. It comes with a data browser, interactive data visualizations, and extensive documentation. The functional API consists of powerful main functions (e.g., `ft_preprocessing`, `ft_freqanalysis, ft_connectivityanalysis`) and a number of smaller auxiliary functions. Most functions can be called with the input data and a `config` structure as input parameters, and return an output data structure that includes a copy of the config, serving as a history of the operations applied to the data and a way to re-apply the analysis to different input data.

EEGLab has been developed since at least 2004 and is an interactive Matlab toolbox for processing continuous and event-related EEG, MEG and other electrophysiological data. It includes both a graphic user interface (GUI) and an API, and has support for user-contributed code via a plug-in interface. Features include interactive visualization, artifact removal, independent component analysis (ICA), time-frequency analysis and source modeling.

The Brainstorm software package is written in Matlab and Java, but can be run as a standalone application without the need for a Matlab license. It focuses on a sophisticated GUI and provides some batch processing functionalities.

Elephant is a Python library for the analysis of electrophysiological data with a focus on generic analysis functions for spike-train data and time-series recordings from electrodes.

MNE Python is a Python package that supports data preprocessing, source localization, statistical analysis, and estimation of functional connectivity between distributed brain regions. It is built on top of the scientific Python ecosystem, has many contributors and is well integrated with other applications using the Neuromag FIF file format. MNE has extensive plotting capabilities and documentation, including publicly available example datasets and tutorials. It supports parallelization on multiple cores of a single machine via Python’s `joblib` module, but currently no direct parallelization support for HPC systems. The API is a combination of fine-grained functions and methods defined directly on the data objects. MNE is focused on the analysis of EEG and MEG data and local field potentials (LFPs) and supports artifact removal, time/frequency analysis and source modeling.

We developed SyNCoPy to complement some of MNE’s and Elephant’s features, offer an easy, FieldTrip-like API, support for time-discrete spike datasets and built-in parallelization on HPC systems.

### The SyNCoPy Architecture

The mentioned software solutions are well established and share different features with SyNCoPy. However none of them is made for handling very large datasets, and for distributed computing on HPC systems. SyNCoPy supports this use case through an architecture that supports trial-parallel out-of-core computations. SyNCoPy’s core data structures consist of metadata and a multi-dimensional data array, but the data array is not loaded into memory by default. Instead, when a computation is requested, the data is streamed trial-wise from HDF5 (Hierarchical Data Format 5) containers stored on the hard disk, and results are written back to disk in a similar fashion. Metadata is stored in JavaScript Object Notation (JSON) format. This approach allows for memory-efficient processing of very large datasets with many trials, as well as for easy trial-based parallelization. On a standard computer, trials can be handled sequentially or in parallel using several cores, if enough memory is available, while on HPC or cloud-based systems, parallelization is achieved by having each node handle one trial at a time. This means that large numbers of trials can be processed in parallel using today’s HPC systems.

The internal architecture of SyNCoPy and the recommended setup for running parallel computations on large datasets is depicted in Figure 1. Users connect to a remote JupyterHub instance, for example provided by an institutional High-Performance Computing (HPC) cluster.

**Figure 1.**
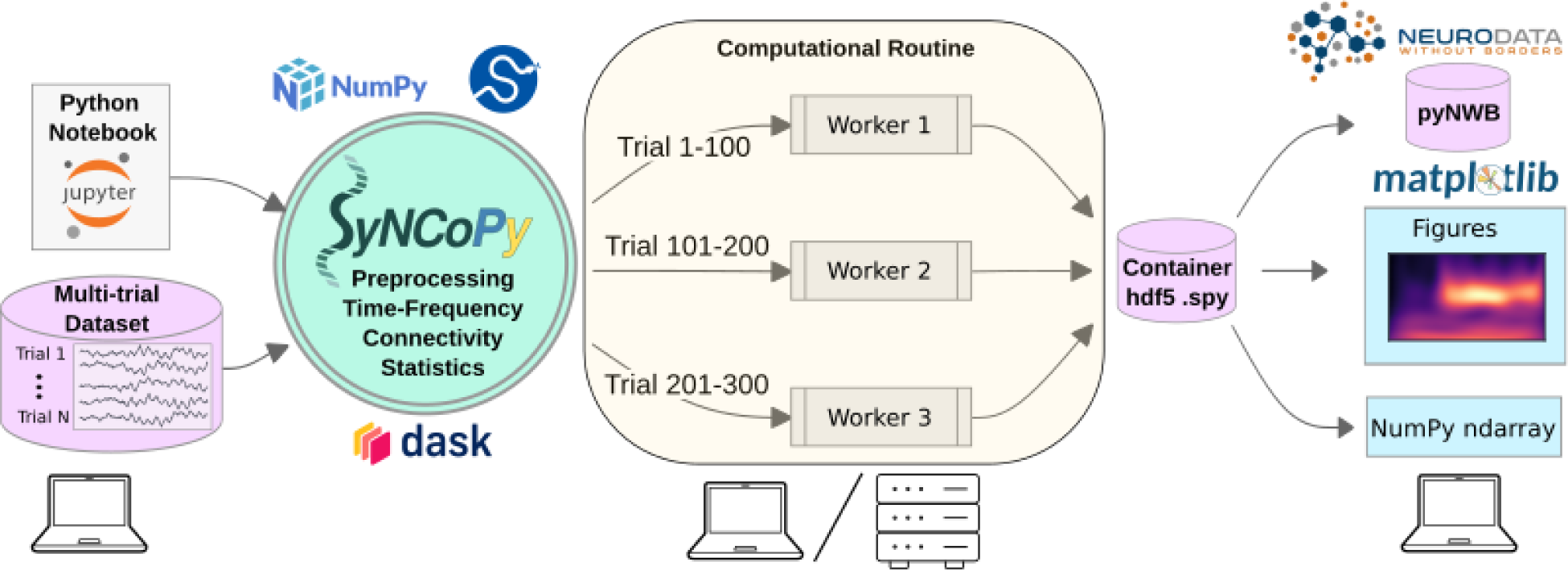
The SyNCoPy architecture and a typical setup for parallel processing. SyNCoPy is started on a laptop (left) to process a multi-trial dataset. When a high-level SyNCoPy API function is executed in a Jupyter Notebook, SyNCoPy’s algorithms based on NumPy and SciPy are wrapped in a computational routine which connects to a high-performance compute cluster (or a local cluster on the laptop) via Dask and automatically distributes the trial-by-trial computations to the available resources. The jobs run in parallel (center), with each worker process handling one job at a time and writing the results for a trial into the proper slot of a single HDF5 container on disk. When all workers have finished their assigned jobs, the results on disk are complete and can be accessed from the SyNCoPy session on the laptop (right). The results can then be visualized with SyNCoPy’s plotting API based on matplotlib, exported to NWB format (Neurodata Without Borders), or NumPy arrays can be extracted directly for custom post-processing using the standard scientific Python tech stack.

After creating a global Dask (Rocklin, 2015) client, running SyNCoPy analyses will use the available computing resources. The input data should reside on fast storage accessible from the cluster, typically a file server. When the user starts a parallel computation, SyNCoPy automatically detects and uses the Dask cluster and distributes the work to the HPC cluster nodes. The nodes write the results to disk, and the SyNCoPy data structure returned by the SyNCoPy API function points to the data on disk. Note that the resulting data is never transferred directly over the network, and is never loaded completely into memory. For post-processing, the API offers interfaces to matplotlib (Hunter, 2007) for plotting, and NumPy (Harris et al., 2020) and pyNWB (Rübel et al., 2022) for data export.

SyNCoPy compute functions (running as a `ComputationalRoutine`) can attach to any running Dask client, and hence harness the full flexibility of the Dask ecosystem, e.g. easy deployment to cloud resources.

SyNCoPy provides specialized data structures and a general method for implementing parallel out-of-core computations on it, the `ComputationalRoutine`. The user-exposed functions (high-level SyNCoPy API, like `syncopy.connectivityanalysis`) internally evaluate user-specified configurations and then use the `ComputationalRoutine` mechanism to execute code that typically works on the data of a single trial. Depending on the global Python environment, the `ComputationalRoutine` executes the per-trial code sequentially, or in parallel via Dask (see also esi-acme^1^) to interact with a parallelization backend, e.g., a Slurm job scheduler running on an HPC cluster. SyNCoPy analysis scripts are agnostic about the hardware environment, meaning analyses can be developed and run locally on single machines like laptops, and the same code can later be deployed on distributed computing resources.

### Feature Overview

The current features of SyNCoPy can be divided into the broad categories data handling, pre-processing, time-locked analysis, frequency-domain analysis, and connectivity-based analysis.

### Data Structures and Data Handling

The data handling category includes functions for loading and saving data using SyNCoPy’s internal data formats, as well as some functions to convert data, i.e. import data and export them into other file formats. SyNCoPy’s core data structures generally contain a multi-dimensional data array and metadata. On disk, the data is represented as an HDF5 file, and when data is loaded into memory, it becomes available as a NumPy array. The data structures can be divided into data types for continuous data and for discrete data. The `AnalogData` class is typically used to store raw electrophysiological data, i.e., multi-channel, regularly-sampled, analog data with one or more trials. If no trial information is available in the data source, the user typically creates a trial definition to define the trials. For many analysis types, latency selections are applied to ensure that the data is time-locked to a certain event like stimulus onset, which results in a `TimeLockData` instance. Algorithms that output real or complex spectral data store these results in instances of the `SpectralData` class, and those resulting in channel-channel interaction information (connectivity measures) return instances of the `CrossSpectralData` class. The discrete data classes `SpikeData` and `EventData` are used to store spikes and events, respectively. The `SpikeData` class can store spikes identified in external spike sorting software like SpyKING Circus (Yger et al., 2018), including the raw waveform around each spike. The `EventData` class is used to store event times, and is typically used in combination with other data classes.

All data classes can be initialized from NumPy arrays and data type-specific metadata, like the sampling frequency for `AnalogData` instances. To facilitate memory-safe data handling also during initialization, Python generators producing single-trial NumPy arrays can be fed directly into the respective SyNCoPy data class constructors. To improve interoperability with other software packages, functions to convert between the data structures of MNE Python and SyNCoPy are available. We also provide functions to save and load data in NWB format (Rübel et al., 2022).

### Pre-Processing

SyNCoPy’s preprocessing functions work on `AnalogData` instances and support detrending, normalizing and filtering signals, including low-pass, high-pass, band-pass and band-stop filters. Resampling and downsampling of time series data is also supported.

### Time-frequency Analysis

SyNCoPy provides functions for frequency analysis and time-frequency analysis on input of type `AnalogData`. The (Multi-)tapered Fourier transform (MTMFFT) algorithms perform spectral analysis on time-series data using either a single taper window or many tapers based on the discrete prolate spheroidal sequence (DPSS). The effective frequency smoothing width can be directly controlled in Hertz with the `tapsmofrq` parameter as in FieldTrip. The single tapers available in SyNCoPy are imported from SciPy’s signal module (Virtanen et al., 2020). The resulting spectra can be post-processed using the FOOOF method (Fitting Oscillations and One-over-f) (Donoghue et al., 2020). A sliding window short-time Fourier transform is also available, as well as Welch’s method for the estimation of power spectra based on time-averaging over short, modified periodograms (Welch, 1967). Both the non-orthogonal continuous wavelet transform (Torrence & Compo, 1998) and superlets, which can reveal fast transient oscillations with high resolution in both time and frequency (Moca et al., 2021), are available in SyNCoPy for time-frequency analysis.

### Connectivity Analysis

The connectivity analysis module reveals functional connectivity between channels. It provides algorithms for cross-spectral density estimation (CSD), coherence, pairwise phase consistency (PPC), (Vinck et al., 2010), nonparametric Granger causality (Dhamala et al., 2008), and cross-correlation. Running connectivity analysis requires `SpectralData` input. If an `AnalogData` instance is passed, an implicit MTFFT analysis is run with default parameters to obtain a `SpectralData` instance.

### Statistics

SyNCoPy provides functions to compute the mean, median, standard deviation, and variance along arbitrary axes of its data classes. The inter-trial coherence can be computed for input of type `SpectralData`. Jackknifing (Richter et al., 2015) is also implemented and can be used to compute confidence intervals for coherence or Granger causality results. The peristimulus time histogram (PSTH) can be computed for `SpikeData` instances (Palm et al., 1988).

### Plotting and Utility Functions

We provide plotting functions for various SyNCoPy data types, including `AnalogData`, `SpectralData` and `SpikeData`. The SyNCoPy plotting functions are intended to give scientists a quick and easy overview of their data during the development of the data analysis pipeline and for project presentations, but not to provide publication-ready figures. The functions internally use matplotlib (Hunter, 2007), and the resulting figures can be post-processed by users if needed.

The `synthdata` module in SyNCoPy contains utility functions to create synthetic datasets, which is useful for training purposes and to test custom algorithms and assess their performance. Apart from standard processes like white noise or Poisson shot noise to simulate spike data, we also offer red noise (AR(1) process) and a phase-diffusion algorithm (Schulze, 2005) to mimic experimental LFP signals.

Basic algebraic operations like addition and multiplication are supported (and parallelized) for all SyNCoPy data classes and NumPy arrays, allowing for flexible synthetic data construction and standard operations like baseline corrections.

### Example Step-by-Step Analysis Pipeline for a Real Electrophysiological Dataset

In the following, we present an example of a step-by-step analysis pipeline to demonstrate how to use SyNCoPy for analyzing extracellular electrophysiology data. For comparison, the same analysis was carried out in Matlab with FieldTrip. The source code for the SyNCoPy version and the FieldTrip version is available online at https://github.com/frieslab/syncopy_paper.

Figure 2 depicts the analysis pipeline and SyNCoPy functions used to process a sample brain signal. The dataset used in the analyses is a publicly available dataset and comes from the Allen Institute Visual Coding – Neuropixels project^2^ and has been described previously (Siegle et al., 2021). In summary, LFP and spiking activity were simultaneously recorded through high-density Neuropixel extracellular electrophysiology probes. These recordings encompass various regions of the mouse brain during the processing of visual stimuli. The LFP data was recorded using Open Ephys (Siegle et al., 2017), and spike data was extracted with Kilosort (Pachitariu et al., 2023). During the experiment, mice were presented with different visual stimuli. Here, the full-field flash stimulus with duration of 250 ms was considered as stimulus epoch, while the 250 ms period before stimulus onset was used as baseline (Figure 2B). In order to also evaluate connectivity analyses, two visual areas from one sample session were selected (Area A, in the Allen dataset referred to as Area VISl, corresponding to Primary visual area, lateral part; Area B, in the Allen dataset referred to as Area VISrl, corresponding to Primary visual area, rostral part). After preprocessing data for aligning the data to stimulus onset, the aforementioned time-domain and frequency-domain analyses were tested on the data. Figure 2C shows the LFP response averaged across different trials and channels of area A. It indicates an evoked response with a short latency after visual stimulus presentation. Time-locked raster plots and the peri-stimulus time histogram (PSTH) of the spike trains are shown for 150 trials of a sample neuron (Figures 2D and 2E, respectively). Subsequently, we calculated the power spectrum of the LFP from Area A, the coherence spectrum between the LFPs of Area A and Area B, the pairwise phase consistency (PPC) spectrum between the LFPs of Area A and Area B, and the nonparametric Granger causality (GC) spectrum between the LFPs of Area A and Area B, as four common frequency-domain analyses.

**Figure 2.**
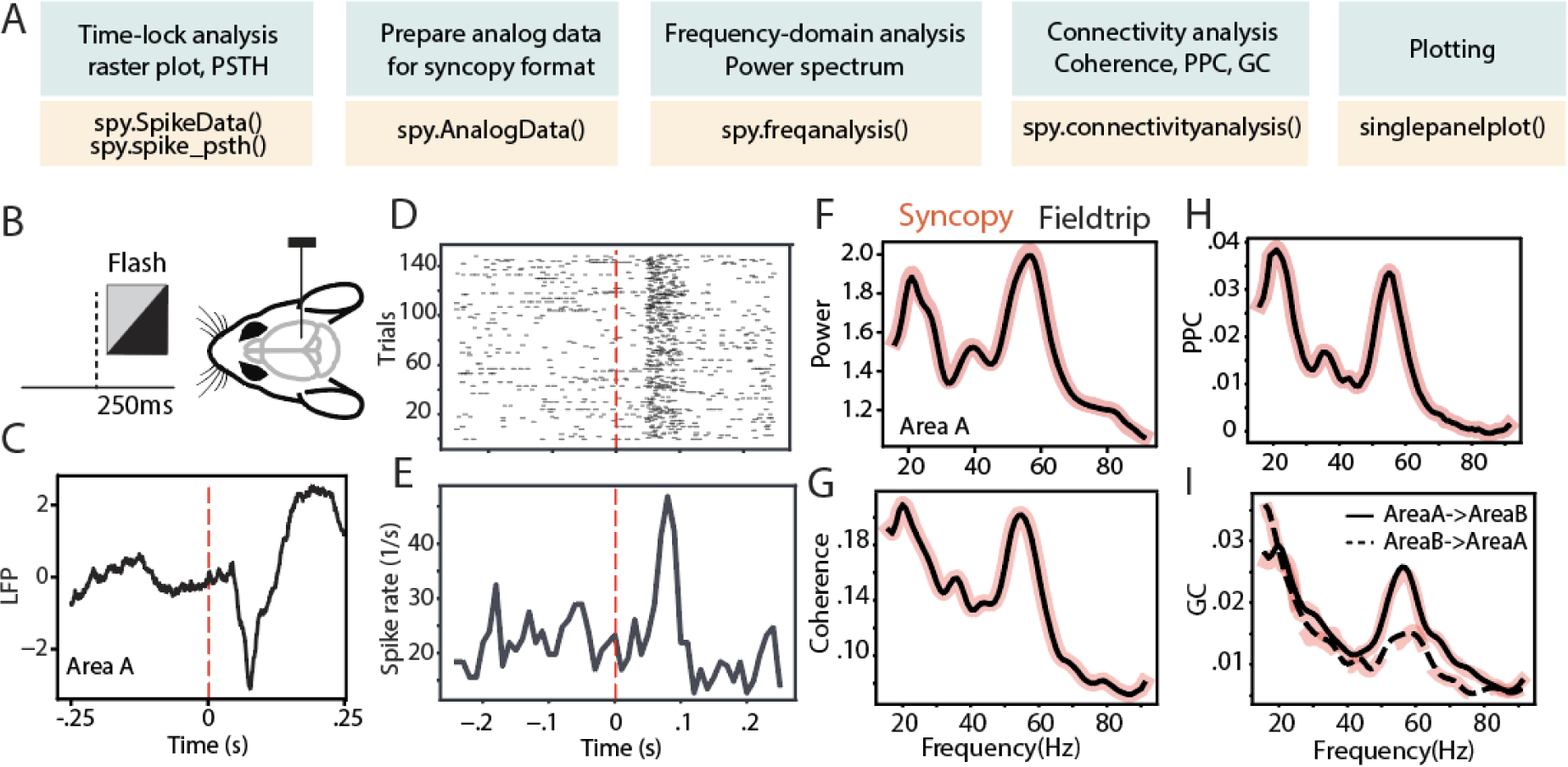
SyNCoPy analysis pipeline for an example electrophysiological dataset. **A)** Analysis pipeline and SyNCoPy functions used to process an example electrophysiological dataset. **B)** During presentation of the full-field flash stimulus lasting for 250 ms, LFP and spiking activity were recorded from different brain areas of awake mice. **C)** The averaged LFP response over trials and channels of area A, time-locked to stimulus onset. **D, E**) Time-lock raster plot (D) and peri-stimulus time histogram (E) of spiking activity of 150 trials in a sample neuron. **F**) Spectra of LFP power ratio between stimulus and baseline period in frequency range of 1-95 Hz averaged over trials and channels of Area A. **G-I**) Same as F but for coherence between LFPs of Area A and Area B (G), pairwise phase consistency between LFPs of Area A and Area B (H), Granger causality between the LFPs of Area A and Area B (I). Black lines are FieldTrip results and red-shaded lines are SyNCoPy results. Solid line is feedforward and dashed line is feedback direction (I).

These analyses were calculated in both SyNCoPy and FieldTrip for demonstration purposes and to illustrate the comparability of the outputs. To this end, the data was first zero padded. Next, based on the MTMFFT method and using the Hann window, the power spectrum was calculated during the stimulus period and the baseline period for each trial and recording channel. MTMFFT conducts frequency analysis on time-series trial data by employing either a single taper (such as Hann) or by utilizing multiple tapers derived from discrete prolate spheroidal sequences (DPSS). For each recording channel separately, the power spectra of the stimulus period and the baseline period were separately averaged, and the ratio of stimulus-over baseline-power calculated. Subsequently, the power-ratio spectra were averaged over channels (Figure 2F).

Similarly, the coherence (Figure 2G), PPC (Figure 2H), and Granger causality (Figure 2I) between the selected area pairs were measured after zero padding the signals. The results are essentially identical between SyNCoPy and FieldTrip for power, coherence, and PPC, and they are very similar for GC (Figures 2F-I).

## Memory benchmarks

### Peak memory consumption - Methods

We investigated the peak memory consumption (PMC) of SyNCoPy for several algorithms in a typical usage scenario, i.e., during parallel processing on an HPC cluster. Specifically, the “small” queue of the Raven cluster at the Max Planck Computing and Data Facility (MPCDF) of the Max-Planck Society was used. In order to assess the memory consumption as a function of the dataset size, we created synthetic datasets of increasing size with SyNCoPy’s `synthdata` module, and processed them with SyNCoPy. We evaluated (1) pre-processing with a Butterworth + Hilbert filter, (2) the MTMFFT, (3) the MTMFFT f.t. algorithm (here, f.t. specifies that a fixed number of tapers was used for better comparison, as explained in more detail below), (4) wavelets, and (5) coherence. The starting dataset size was 10 trials, 5000 samples per trial and 50 channels, which requires about 10 MB of space. We created scripts to run each algorithm with different dataset sizes. After each call to a SyNCoPy API function, the Python garbage collector was called to ensure meaningful measurements. During each run, the PMC was monitored with the `memory_profiler` package^3^ for Python. The PMC is the highest amount of memory consumption of the submitting process and one worker that was measured during a run. We repeated the process 20 times for each unique combination of dataset size and algorithm to obtain robust results. We report the mean and the standard deviation over the 20 runs in Figure 3.

**Figure 3.**
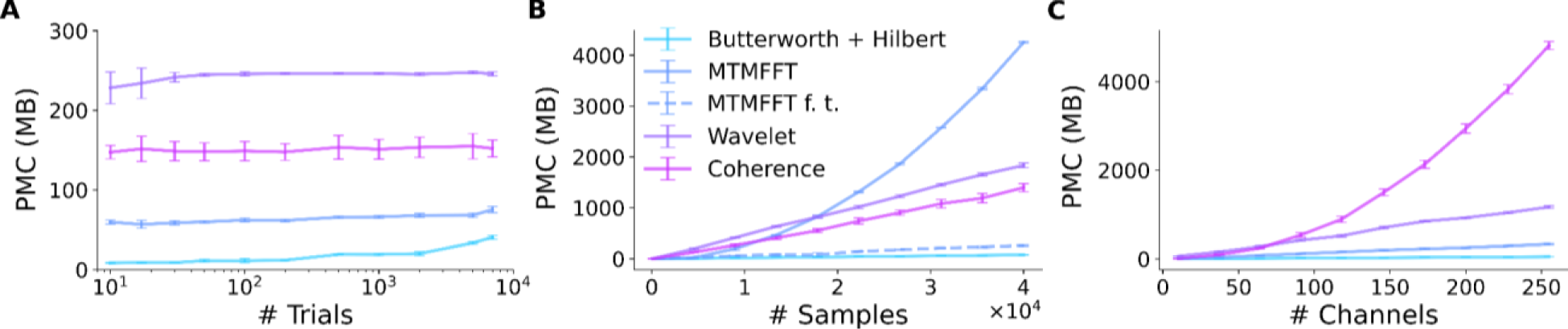
SyNCoPy memory efficiency. Peak memory consumption (PMC) as a function of input size for selected algorithms. The PMC measurements are based on synthetic data. The starting dataset size is 10 trials, 5000 samples and 50 channels. Each datapoint shows PMC mean and standard deviation of 20 independent runs. **A)** PMC is largely independent of the number of trials. The total size of the test dataset varied over almost three orders of magnitude (10 trials to 7,000 trials, ∼10MB to 7GB), while the size of a single trial was kept constant at 1MB. **B)** PMC depends on the number of samples per trial and the algorithm. The number of samples (length of the signals) varied from 10 to 40 000. **C**) PMC depends on the number of channels and the algorithm. Channel numbers varied from two to 250.

### Peak memory consumption - Results

The results of the peak memory consumption (PMC) measurements are illustrated in Figure 3. First, we investigated the effect of the trial count on PMC (Figure 3A). We incrementally increased the number of trials from 10 up to 40,000 while keeping the samples per trial and the number of channels constant. At each datapoint, we performed 20 independent runs with the respective algorithm. The PMC stayed largely constant, irrespective of the trial count, for all algorithms. The PMC was lowest for the Butterworth filter, followed by the MTMFFT, Coherence and Wavelets. Second, we demonstrate the effect of increasing the number of samples per trial on memory consumption (Figure 3B). We gradually increased the number of samples per trial from 10 to 40,000 while keeping the trial count and channel count constant. For the wavelets, the multi-taper analysis with a fixed number of tapers (MTMFFT f.t.), and coherence computation, a linear effect on the PMC is visible. For the Butterworth filter, memory consumption is essentially constant, as for this method we employ SciPy’s `signal.sosfiltfilt` implementation, which works on finite sections of the input data. For the full multi-taper analysis (MTMFFT), the PCM increases quadratically with the sample count: The FFT itself has a PMC that is a linear function of the number of samples, and the number of tapers needed to achieve a consistent frequency smoothing (tapsmofrq parameter) also scales with the number of samples. Finally, we observe the effect of increasing the number of channels on the PMC of the algorithms (Figure 3C). For the wavelets and the MTMFFT, a linear effect on the PMC is shown. For the Butterworth filter, the PMC again is almost constant. Coherence shows quadratic scaling of PMC with the number of channels, which directly follows from combinatorics

## Discussion

SyNCoPy is a Python package for the analysis of electrophysiological data, with a focus on extracellular electrophysiology. It stands out from similar software packages by its ability to scale easily from laptops to HPC systems and thus support very large datasets, and an API similar to FieldTrip’s. SyNCoPy’s support for big data is based on its architecture, which (1) allows for easy usage of typical HPC systems available at many scientific institutions, (2) streams data from disk to memory only when needed and (3) isolates computations on the minimal amount of data required for independent computations. We demonstrated SyNCoPy’s memory efficiency by benchmarking peak memory consumption (PMC) for a number of algorithms. The results demonstrate that SyNCoPy’s architecture is indeed able to provide largely constant PMC, independent of the number of trials. Moreover, the PMC scales as expected for the respective algorithms with increases in single-trial size.

From a feature perspective, SyNCoPy currently focuses on preprocessing of raw data, time-frequency analysis and connectivity measures. We expect that neuroscience users may want to employ SyNCoPy in combination with other well-established software packages like MNE Python, Elephant (Denker et al., 2023) and others that contain complementary functionality. To facilitate this, we provide support for converting MNE Python data structures and importing and exporting standard file formats like NWB. Also, the SyNCoPy file format is based on the open standards HDF5 and JSON, and can thus be read by standard libraries available for a variety of languages.

SyNCoPy does not have a graphical user interface and relies on scripting. While this may require a certain initial time investment for users completely new to programming, we believe that the standardization and increased reproducibility offered by this approach pays off quickly. FieldTrip is largely based on the same approach and has reached a large user community. To help new users, SyNCoPy comes with full API documentation and includes a set of articles that demonstrate typical analysis workflows. Questions and issues can be reported and discussed on the SyNCoPy Github repository^4^.

### Limitations

First, it is important to acknowledge that memory efficiency is a software requirement that, in some situations, conflicts with performance in the sense of processing speed: for a small dataset, it is faster to load everything into memory at once than to stream chunks of the data on demand. However, for large datasets, this computing strategy prevents the processing of datasets larger than (a certain fraction of) the machine’s RAM and thus is not feasible.

Second, SyNCoPy is focused on trial-parallel processing, which is from our perspective a very common scenario in Neuroscience. However, in some situations or for certain algorithms, it may be beneficial to support parallelization along different axes. While SyNCoPy does have built-in support for parallelization over channels for some algorithms, it does not in general support parallelization along an arbitrary axis of the data set.

Third, extension of SyNCoPy with new algorithms is possible by creating a custom Computational Routine, but this process currently requires a good understanding of both parallel computing and some SyNCoPy internals, and is thus intended for more advanced users.

Fourth, the target audience of SyNCoPy consists of neuroscientists who need to process larger datasets. The exact limitation for the size of the data set depends on the specific algorithms and the settings used, of course. But what always holds is that a single trial must easily fit into the RAM of the machine, i.e., typically the HPC cluster node that runs the computations. It is important to understand that certain operations used while loading and saving data, or in the algorithms themselves, will need to create one, or in some cases even more, copies of the trial data in memory. Therefore, working with a dataset that has almost the size of the RAM is not feasible in reality. This is not a limitation of SyNCoPy, but applies to all operations on computers, including the standard NumPy and SciPy libraries used internally by SyNCoPy to implement or run the algorithms on the data of a single trial. The required memory typically is a small multiple of the single-trial size.

### Conclusion

SyNCoPy provides seamless scaling of trial-based workflows for the analysis of large electrophysiology datasets in Python. In this paper, we demonstrated its ability to scale to very large datasets by measuring the peak memory consumption over a range of algorithms for data sets with varying numbers of trials, samples per trial, and channels. Furthermore, we illustrated how to use SyNCoPy on a real-world dataset, along with a direct comparison of the same analyses carried out with the well-established FieldTrip toolbox.

SyNCoPy was built to integrate well into the current ecosystem of neuroscience tools. We hope that it will help researchers to work with large datasets in a reproducible way and lower the barrier to fully utilize existing HPC resources in neuroscience.

### Availability

SyNCoPy is free software, available at https://github.com/esi-neuroscience/syncopy and on PyPI and conda-forge.

## Acknowledgements

The authors would like to thank Robert Oostenveld and Jan-Mathijs Schoffelen for their invaluable input and insightful feedback throughout the development of SyNCoPy, Mukesh Dhamala for providing help with the Python implementation of the Granger causality algorithm, and Katharine Shapcott and Muad Abd El Hay for early testing of the software, suggestions and discussions. Additionally, the authors acknowledge the Max Planck Computing and Data Facility (MPCDF) and the IT team at ESI for providing generous access to high-performance computing resources.

## Author Contributions

GM: Core development, focus on methods and algorithms. Memory benchmarking. TS: Core development, focus on backend engineering and visualizations. Wrote the initial draft of the manuscript. DSK: Software testing, contributions to software design and development. MP: Data analysis SyNCoPy and FieldTrip. SF: Core development, focus on data backend. JS: Contribution to core development. PF: Initiated and conceptualized the project, prioritization of software functionality, input from neuroscience perspective, funding. All authors read and approved the final version of the manuscript.

## Footnotes

1 https://github.com/esi-neuroscience/acme

2 https://allensdk.readthedocs.io/en/latest/visual_coding_neuropixels.html

3 https://github.com/pythonprofilers/memory_profiler

4 https://github.com/esi-neuroscience/syncopy/issues

## References

1. Delorme, A., & Makeig, S. (2004). EEGLAB: An open source toolbox for analysis of single-trial EEG dynamics including independent component analysis. Journal of Neuroscience Methods, 134(1), 9–21. 10.1016/j.jneumeth.2003.10.009

2. Delorme, A., Mullen, T., Kothe, C., Akalin Acar, Z., Bigdely-Shamlo, N., Vankov, A., & Makeig, S. (2011). EEGLAB, SIFT, NFT, BCILAB, and ERICA: New Tools for Advanced EEG Processing. Computational Intelligence and Neuroscience, 2011, e130714. 10.1155/2011/130714

3. Denker, M., Köhler, C., Jurkus, R., Kern, M., Kurth, A. C., Kleinjohann, A., Bouss, P., Davison, A., Morales-Gregorio, A., Kramer, M., & Ito, J. (2023). *Elephant 0.13.0* [Computer software]. Zenodo. 10.5281/zenodo.8144467

4. Dhamala, M., Rangarajan, G., & Ding, M. (2008). Analyzing information flow in brain networks with nonparametric Granger causality. NeuroImage, 41(2), 354–362. 10.1016/j.neuroimage.2008.02.020

5. Donoghue, T., Haller, M., Peterson, E. J., Varma, P., Sebastian, P., Gao, R., Noto, T., Lara, A. H., Wallis, J. D., Knight, R. T., Shestyuk, A., & Voytek, B. (2020). Parameterizing neural power spectra into periodic and aperiodic components. Nature Neuroscience, 23(12), Article 12. 10.1038/s41593-020-00744-x

6. Gramfort, A., Luessi, M., Larson, E., Engemann, D. A., Strohmeier, D., Brodbeck, C., Parkkonen, L., & Hämäläinen, M. S. (2014). MNE software for processing MEG and EEG data. NeuroImage, 86, 446–460. 10.1016/j.neuroimage.2013.10.027

7. Gramfort, A., Luessi, M., Larson, E., Engemann, D., Strohmeier, D., Brodbeck, C., Goj, R., Jas, M., Brooks, T., Parkkonen, L., & Hämäläinen, M. (2013). MEG and EEG data analysis with MNE-Python. Frontiers in Neuroscience, 7. https://www.frontiersin.org/articles/10.3389/fnins.2013.00267

8. Harris, C. R., Millman, K. J., van der Walt, S. J., Gommers, R., Virtanen, P., Cournapeau, D., Wieser, E., Taylor, J., Berg, S., Smith, N. J., Kern, R., Picus, M., Hoyer, S., van Kerkwijk, M. H., Brett, M., Haldane, A., del Río, J. F., Wiebe, M., Peterson, P.,…Oliphant, T. E. (2020). Array programming with NumPy. Nature, 585(7825), Article 7825. 10.1038/s41586-020-2649-2

9. Hunter, J. D. (2007). Matplotlib: A 2D Graphics Environment. Computing in Science & Engineering, 9(3), 90–95. 10.1109/MCSE.2007.55

10. Moca, V. V., Bârzan, H., Nagy-Dăbâcan, A., & Mureșan, R. C. (2021). Time-frequency super-resolution with superlets. Nature Communications, 12(1), Article 1. 10.1038/s41467-020-20539-9

11. Oostenveld, R., Fries, P., Maris, E., & Schoffelen, J.-M. (2011). FieldTrip: Open source software for advanced analysis of MEG, EEG, and invasive electrophysiological data. Computational Intelligence and Neuroscience, 2011, 1:1–1:9. 10.1155/2011/156869

12. Pachitariu, M., Sridhar, S., & Stringer, C. (2023). Solving the spike sorting problem with Kilosort (p. 2023.01.07.523036). bioRxiv. 10.1101/2023.01.07.523036

13. Palm, G., Aertsen, A. M. H. J., & Gerstein, G. L. (1988). On the significance of correlations among neuronal spike trains. Biological Cybernetics, 59(1), 1–11. 10.1007/BF00336885

14. Richter, C. G., Thompson, W. H., Bosman, C. A., & Fries, P. (2015). A jackknife approach to quantifying single-trial correlation between covariance-based metrics undefined on a single-trial basis. NeuroImage, 114, 57–70. 10.1016/j.neuroimage.2015.04.040

15. Rübel, O., Tritt, A., Ly, R., Dichter, B. K., Ghosh, S., Niu, L., Baker, P., Soltesz, I., Ng, L., Svoboda, K., Frank, L., & Bouchard, K. E. (2022). The Neurodata Without Borders ecosystem for neurophysiological data science. eLife, 11, e78362. 10.7554/eLife.78362

16. Schulze, H. (2005). Stochastic Models for Phase Noise. In International OFDM-Workshop.

17. Siegle, J. H., Jia, X., Durand, S., Gale, S., Bennett, C., Graddis, N., Heller, G., Ramirez, T. K., Choi, H., Luviano, J. A., Groblewski, P. A., Ahmed, R., Arkhipov, A., Bernard, A., Billeh, Y. N., Brown, D., Buice, M. A., Cain, N., Caldejon, S.,…Koch, C. (2021). Survey of spiking in the mouse visual system reveals functional hierarchy. Nature, 592(7852), Article 7852. 10.1038/s41586-020-03171-x

18. Siegle, J. H., López, A. C., Patel, Y. A., Abramov, K., Ohayon, S., & Voigts, J. (2017). Open Ephys: An open-source, plugin-based platform for multichannel electrophysiology. Journal of Neural Engineering, 14(4), 045003. 10.1088/1741-2552/aa5eea

19. Tadel, F., Baillet, S., Mosher, J. C., Pantazis, D., & Leahy, R. M. (2011). Brainstorm: A User-Friendly Application for MEG/EEG Analysis. Computational Intelligence and Neuroscience, 2011, e879716. 10.1155/2011/879716

20. Torrence, C., & Compo, G. P. (1998). A Practical Guide to Wavelet Analysis. Bulletin of the American Meteorological Society, 79(1), 61–78. 10.1175/1520-0477(1998)079<0061:APGTWA>2.0.CO;2

21. Vinck, M., van Wingerden, M., Womelsdorf, T., Fries, P., & Pennartz, C. M. A. (2010). The pairwise phase consistency: A bias-free measure of rhythmic neuronal synchronization. NeuroImage, 51(1), 112–122. 10.1016/j.neuroimage.2010.01.073

22. Virtanen, P., Gommers, R., Oliphant, T. E., Haberland, M., Reddy, T., Cournapeau, D., Burovski, E., Peterson, P., Weckesser, W., Bright, J., van der Walt, S. J., Brett, M., Wilson, J., Millman, K. J., Mayorov, N., Nelson, A. R. J., Jones, E., Kern, R., Larson, E.,…van Mulbregt, P. (2020). SciPy 1.0: Fundamental algorithms for scientific computing in Python. Nature Methods, 17(3), Article 3. 10.1038/s41592-019-0686-2

23. Welch, P. (1967). The use of fast Fourier transform for the estimation of power spectra: A method based on time averaging over short, modified periodograms. IEEE Transactions on Audio and Electroacoustics, 15(2), 70–73. 10.1109/TAU.1967.1161901

24. Yger, P., Spampinato, G. L., Esposito, E., Lefebvre, B., Deny, S., Gardella, C., Stimberg, M., Jetter, F., Zeck, G., Picaud, S., Duebel, J., & Marre, O. (2018). A spike sorting toolbox for up to thousands of electrodes validated with ground truth recordings in vitro and in vivo. eLife, 7, e34518. 10.7554/eLife.34518

